# An amino acid motif in HLA-DRβ1 distinguishes patients with uveitis in juvenile idiopathic arthritis

**DOI:** 10.1101/140954

**Authors:** A.J.W. Haasnoot, M.W. Schilham, S.S.M. Kamphuis, P.C.E. Hissink Muller, A. Heiligenhaus, D. Foell, R.A. Ophoff, T.R.D.J. Radstake, A.I. Den Hollander, T.H.C.M. Reinards, S. Hiddingh, N. Schalij-Delfos, E.P.A.H. Hoppenreijs, M.A.J. van Rossum, C. Wouters, R.K. Saurenmann, N. Wulffraat, ICON-JIA study group, R. ten Cate, J.H. de Boer, S.L. Pulit, J.J.W. Kuiper

## Abstract

Uveitis is a visually-debilitating disorder that affects up to 30% of children with juvenile idiopathic arthritis (JIA). To identify genetic susceptibility loci for uveitis in JIA, we conducted a genome-wide association study comparing 192 JIA-associated uveitis cases with 330 JIA individuals without uveitis. Two cohorts of JIA patients underwent genotyping and quality control. We used an HLA-specific imputation panel to impute HLA-specific amino acids and HLA types, and identified the amino acid serine at position 11 (serine-11) in *HLA- DRB1* as associated to increased risk of uveitis (OR = 2.60, p = 5.43×10^−10^). The signal at serine-11 was female-specific (interaction of sex and serine-11, p = 0.0096). Serine-11 resides in the YST-motif (positions 10-12) in the peptide binding groove of HLA-DRB1. Quantitative binding affinity predictions revealed peptide-binding preferences that distinguish HLA-DRB1 allotypes with the YST-motif. Our findings highlight a genetically distinct, sexually-dimorphic feature of JIA-associated uveitis.

## Introduction

Juvenile idiopathic arthritis (JIA) is the most common chronic rheumatic disease in childhood,^1,2^ affecting approximately 16-150 per 100,000 individuals.^3^ As many as 1 in 3 children that suffer from the most common JIA categories, oligoarticular and polyarticular rheumatoid factor-negative JIA, develop uveitis, a chronic inflammatory eye disease, which is the most frequent extra-articular manifestation of JIA.^4,5^ Uveitis threatens the sight of those affected, and can result in complications including macular edema, band keratopathy, cataracts, glaucoma, and hypotony.^4^ Because it typically afflicts the young (< 7 years old), uveitis dramatically impacts quality of life both in childhood and as an adult.^4,6^

Early detection and adequate ophthalmological management of JIA-associated uveitis are critical to prevent significant, sight-threatening complications.^7–9^ Despite its severity, uveitis is typically insidious in onset and often becomes symptomatic only after irreversible damage has occurred. Consequently, rigorous ophthalmological screening of all JIA patients is required for early detection and treatment of uveitis associated with JIA.^10^ Nonetheless, severe ocular complications may already be present at the time of a uveitis diagnosis. Several clinical factors are associated with increased uveitis risk: young age (< 7 years old) at JIA onset; the presence of antinuclear antibodies (ANA), elevated erythrocyte sedimentation rate (ESR) at JIA onset, and JIA category (oligoarticular and polyarticular rheumatoid factor (RF)-negative).^10–12^ Sex is also considered a risk factor, though the available epidemiologic data is controversial. Female patients comprise the majority of JIA categories with uveitis, and are therefore observed somewhat more frequently (but only modestly so) in the overall JIA population. The extent to which biological risk for JIA-associated uveitis is sexually dimorphic, however, is not known.^12^

Both JIA and JIA-associated uveitis (i.e., JIA-uveitis) are multifactorial autoimmune disorders; each has a strong, though not fully penetrant, genetic predisposition.^4,13^ Both conditions are complex and driven by biological and environmental factors.^4,13^ There have been no reports of observed Mendelian inheritance patterns of uveitis in JIA-affected families,^4^ further indicating a complex trait architecture. Genome-wide association studies in JIA individuals (with and without uveitis) have revealed a number of loci associated to the disease,^14–16^ including multiple independent loci in the major histocompatibility complex (MHC) which collectively explain >20% of the phenotypic variation in broadly-defined JIA.^16^ A number of HLA types have also been identified as increasing risk of JIA-associated uveitis compared to controls.^4,17–19^ However, genome-wide genetic markers that distinguish JIA without uveitis from JIA-associated uveitis remain elusive, and the exact disease mechanisms predisposing only a subgroup of children to developing uveitis are unknown. Due to the severity of uveitis and the complications surrounding its detection, biomarkers to predict the development of uveitis would help in early detection of the presence of JIA-uveitis in high-risk children without frequent, extensive, and costly screenings for all JIA patients.

Here, we performed genotyping in 192 JIA-associated uveitis cases and 330 JIA patients without uveitis for at least 4 years, with the aim of identifying those genetic variants that segregate more commonly in JIA patients with uveitis compared to those without uveitis. All samples analysed here fall into the following *International League of Associations for Rheumatology* (ILAR) categories: (i) oligoarticular extended, (ii) oligoarticular persistent, and (iii) RF-negative polyarticular JIA. We analyzed the collective JIA-uveitis categories as one group to increase power, because these common categories of JIA are all particularly prone to uveitis and have recently been shown to be genetically highly similar in HLA associations.^20^ To identify risk variants in a high-density set of polymorphisms, we performed two sets of imputation: one using a sequencing-based reference panel, to scan for risk-increasing alleles genome-wide; and a second, with an MHC-specific reference panel, to identify additional SNPs, amino acids, and HLA types that confer risk to disease. We identified serine (S) at position 11 in *HLA-DRB1* as highly associated to increased risk of uveitis (odds ratio (OR) = 2.60 [95% CI: 1.92 - 3.52], p = 5.46×10^−10^). Sex-specific analysis revealed that the association at serine-11 in *HLA-DRB1* was specific to females only (p_females_ = 7.61×10^−10^, p_males_ = 0.18). Quantitative prediction of peptide binding revealed that the presence of serine-11, which is located in the peptide-binding groove, affects overall peptide-binding preferences of common HLA-DRB1 allotypes.

## Results

We performed our analysis in two phases that we then jointly analyzed to improve power for locus discovery.^21^ In Phase 1, we genotyped 137 JIA-associated uveitis cases, 247 non-uveitis JIA samples, and 398 population-level controls without JIA or uveitis, all collected from within the Netherlands. Samples were genotyped on the HumanOmniExpress24 array, containing ∼700,000 markers. Samples and SNPs underwent quality control (QC) (**Materials and Methods**) to remove those samples and SNPs with low-quality genotyping data. After QC, 126 JIA-associated uveitis cases, 231 non-uveitis JIA samples, and 394 population-level controls remained (Table 1). We prephased the genotyping data using SHAPEIT2^22–24^ and imputed the phased haplotypes with IMPUTE2,^25–27^ using a reference panel constructed from the 2,504 global samples included in the 1000 Genomes Project Phase 3.^28^ In parallel to genome-wide imputation, we extracted the MHC from the genotype data and imputed additional SNPs, amino acids and HLA types using the SNP2HLA pipeline^29^ (**Materials and Methods**).

**Table 1.** Samples included in Phase 1 and Phase 2 of the analysis. Sample numbers are shown both pre- and post-quality control (QC) for JIA cases with (JIA-uveitis) and without uveitis (JIA non-uveitis). The proportion of patients that are positive for antinuclear antibodies (ANA+) as well as sex distribution are shown.

|  | JIA patients, total | JIA-uveitis (%) | JIA non-uveitis (%) |
| --- | --- | --- | --- |
| *Number of samples pre-QC* |  |  |  |
| Phase 1 | 384 | 137 (36) | 247 (64) |
| Phase 2 | 192 | 77 (40) | 115 (60) |
| Total | **576** | **214 (37)** | **362 (63)** |
| *Number of samples Post-QC* |  |  |  |
| Phase 1 | 357 | 126 (35) | 231 (65) |
| Phase 2 | 165 | 66 (40) | 99 (60) |
| Total | **522** | **192 (37)** | **330 (63)** |
| ANA-status: |  |  |  |
| ANA+ | 222* | 117 (53) | 105 (47) |
| ANA- | 134* | 25 (19) | 109 (81) |
| Sex: |  |  |  |
| Female | 364 | 136 (37) | 228 (63) |
| Male | 158 | 56 (65) | 102 (35) |

*\*total number of cases with available data*

In Phase 2, an additional 77 JIA-associated uveitis cases and 115 JIA without uveitis samples (total N = 192), collected from Germany, Belgium and Switzerland, were genotyped on the same array. Samples and SNPs underwent QC nearly identical to that of Phase 1 (**Materials and Methods**). After QC, 66 JIA-associated uveitis cases and 99 non-uveitis JIA samples remained for analysis. The cleaned genotyping data was prephased and imputed both genome-wide and within the MHC, precisely as in Phase 1.

### Genome-wide association testing

For genome-wide association testing in the Phase 1 and Phase 2 data, we kept those SNPs with a minor allele frequency (MAF) > 1% and with an imputation quality score > 0.7; >9.2M variants were available in both the Phase 1 and Phase 2 after imputation and filtering (82.6% and 77.6% of SNPs in Phase 1 and Phase 2 respectively had an info score ≥ 0.9; **Supplementary Table 1**). For MHC association testing, we analyzed only those variants with MAF > 1%. No imputation quality filter was applied to the MHC-specific imputed data; 97.7% and 96.4% of MHC variants in Phase 1 and Phase 2, respectively, had an imputation quality score > 0.9 (**Supplementary Table 1**).

We performed genome-wide association testing across the autosomal genome, using a logistic regression framework assuming an additive model in PLINK 1.9^30,31^, correcting for sex and the first two principal components (**Supplementary Figures 1 and 2**). Our data enabled three possible comparison groups: (a) JIA cases and population-level controls, (b) JIA-associated uveitis cases and population-level controls, and (c) JIA-associated uveitis cases and non-uveitis JIA. As the goal of our analysis was to investigate similarities and differences in the genetic architecture of JIA and uveitis, and because far larger GWAS of JIA and population-level controls have already been performed,^20^ we performed the first two genome-wide association studies (GWAS) as a means of data quality control only (**Supplementary Figure 3**) and account for them when considering multiple testing burden. We focus here on the results of the last comparison: JIA-associated uveitis cases and non-uveitis JIA samples.

We performed separate GWAS in Phase 1 and Phase 2 and then combined the summary-level results in an inverse variance-weighted fixed effects meta-analysis in METAL,^32^ for a combined analysis of 330 non-uveitis JIA samples and 192 uveitis cases. Combining data, rather than performing a separate discovery and replication phase, improves power for discovery of novel association signals.^21^ In the genome-wide scan, the single signal achieving genome-wide significance (set at p < 2×10^−8^, after adjusting for three phenotype comparisons) resided in the MHC (**Supplementary Figures 1 and 2**).

### Association testing and fine-mapping in the MHC

To identify the amino acids or HLA types driving the genome-wide association signal, we performed a mega-analysis across the imputed MHC data. We first merged imputation dosages from Phase 1 and Phase 2, and then used PLINK 1.9^31^ to perform logistic regression, assuming an additive model and correcting for the top 5 principal components, sex, and analysis phase (i.e., Phase 1 or Phase 2). This ‘mega-analysis’ approach is theoretically and empirically highly similar to inverse variance-weighted meta-analysis.^33,34^ We also performed a meta-analysis across Phase 1 and Phase 2 and found that, indeed, the odds ratios derived from mega-analysis and meta-analysis in the MHC were highly concordant (Pearson’s r = 0.95, **Supplementary Figure 4**). The mega-analysis allows for the additional advantage of allowing for interaction testing and conditional analysis on any associated variants across the full dataset.

Of the SNPs, amino acids, and classical alleles tested in the MHC, we observed the strongest association for the presence of either serine (S) or aspartic acid (D) at position 11 in *HLA-DRB1* (OR = 2.59 [95% CI: 1.92 - 3.50], p = 4.80×10^−10^; Figure 1, Table 2). To identify which of the possible residues at position 11 explain the signal, we first performed an omnibus test of all but one of the 6 alleles present at *HLA-DRB1* position 11 compared to a null model. We found that the goodness-of-fit of the omnibus test far exceed that of a null model including only sex, principal components and genotyping batch (likelihood ratio test p = 1.5×10^−9^). To test if a single *HLA-DRB1* position 11 allele was driving the top association signal (as opposed to all possible alleles together, as modeled by the omnibus test), we conditioned the top association signal first on serine and then on aspartic acid at position 11, by including the dosages of the respective variant as an additional covariate in the logistic regression model (Figure 1, Table 2, **Supplementary Table 2**). After conditioning on serine at position 11, the association signal at position 11 dropped substantially (p = 0.069), while conditioning on aspartic acid at position 11 left the association signal essentially unchanged (OR = 2.60 [95% CI: 1.92 - 3.52], p = 5.43×10^−10^), indicating that serine explains the bulk of the signal. The association at serine-11 in *HLA-DRB1* was consistent in direction of effect and magnitude of association in both Phase 1 (OR = 2.15 [95% CI: 1.52 - 3.05], p = 1.59×10^−5^) and Phase 2 (OR = 3.34 [95% CI: 1.92 - 5.83], p = 2.11×10^−5^), indicating that both phases of the study contributed to the joint association signal^21^ in the mega-analysis.

**Table 2.** Association results for amino acids in *HLA-DRB1*. The top results from the mega-analysis in uveitis and JIA without uveitis in *HLA-DRB1*. All reported results correspond to presence of the given amino acid residue(s). Presence of serine (Ser) or aspartic acid (Asp) at position 11 in *HLA-DRB1* (imputation info score = 1.11) was the top hit after initial association testing. Conditioning on either aspartic acid or serine revealed that the amino acids at positions 10-13, all well-imputed (imputation info = 1.08) and in perfect linkage disequilibrium, explain the bulk of the initial signal.

| *DRβ1*<br>position | Amino acid<br>residue(s) | Classical HLA<br>alleles | Samples<br>(all, females, males) | Mega-analysis |  |  |
| --- | --- | --- | --- | --- | --- | --- |
|  |  |  |  | Freq.<br>Case \| Cntl | OR [95% CI] | P-value |
| Initial association testing |  |  |  |  |  |  |
| 11 | Ser or Asp | *03,*08,*09,<br>*11,*12,*13,*14 | All | 0.79 \| 0.59 | 2.59 [1.92 - 3.50] | 4.80 × 10<sup>-10</sup> |
| 11 | Ser | *03,*08,*11,<br>*12,*13,*14 | All | 0.77 \| 0.57 | 2.47 [1.84 - 3.32] | 1.44 × 10<sup>-9</sup> |
| Conditioning on aspartic acid (Asp) at position 11 |  |  |  |  |  |  |
| 11 | Ser or Asp | *03,*08,*09,<br>*11,*12,*13,*14 | All | 0.79 \| 0.59 | 2.60 [1.92 - 3.52] | 5.43 × 10<sup>-10</sup> |
| 11 | Ser | *03,*08,*11,<br>*12,*13,*14 |  | 0.77 \| 0.57 | 2.60 [1.92 - 3.52] | 5.46 × 10<sup>-10</sup> |
| Conditioning on serine (Ser) at position 11 |  |  |  |  |  |  |
| 11 | Ser or Asp | *03,*08,*09,<br>*11,*12,*13,*14 | All | 0.79 \| 0.59 | 2.36 [0.93 - 5.98] | 0.069 |
| Sex-specific association testing |  |  |  |  |  |  |
| 233 | Thr | *01,*04,*07,*08,<br>*09,*10,*15, *16 | Females | 0.18 \| 0.44 | 0.30 [0.20 - 0.43] | 3.50 × 10<sup>-10</sup> |
|  |  |  | Males | 0.37 \| 0.46 | 0.73 [0.44 - 1.20] | 0.210 |
| 11 | Ser or Asp | *03,*08,*09,<br>*11,*12,*13,*14 | Females | 0.84 \| 0.59 | 3.48 [2.35 - 5.16] | 4.92 × 10<sup>-10</sup> |
|  |  |  | Males | 0.67 \| 0.56 | 1.52 [0.92 - 2.51] | 0.100 |
| 11 | Ser | *03,*08,*11, | Females | 0.82 \| 0.57 | 3.30 [2.26 - 4.83] | 7.61 × 10<sup>-10</sup> |
|  |  | *12,*13,*14 | Males | 0.64 \| 0.55 | 1.41 [0.86 - 2.31] | 0.177 |
| Conditioning on threonine (Thr) at position 233 |  |  |  |  |  |  |
| 11 | Ser or Asp | *03,*08,*09,<br>*11,*12,*13,*14 | Females | 0.84 \| 0.59 | 1.98 [0.73 - 5.32] | 0.178 |
|  |  |  | Males | 0.67 \| 0.56 | 3.14 [0.70 - 14.15] | 0.136 |
| 11 | Ser | *03,*08,*11,<br>*12,*13,*14 | Females | 0.82 \| 0.57 | 0.84 [0.10 - 6.94] | 0.878 |
|  |  |  | Males | 0.64 \| 0.55 | 2.83 [0.16 - 51.47] | 0.482 |
| Sex-interaction association testing |  |  |  |  |  |  |
| 11 | Ser*Sex | *03,*08,*11,<br>*12,*13,*14 | All | -- | 2.25 [1.22 - 4.15] | 0.0096 |

Association statistics for tyrosine (position 10), threonine (position 12) and serine or glycine (position 13) are identical to the association statistics reported for serine (position 11) due to linkage disequilibrium.

**Figure 1.**
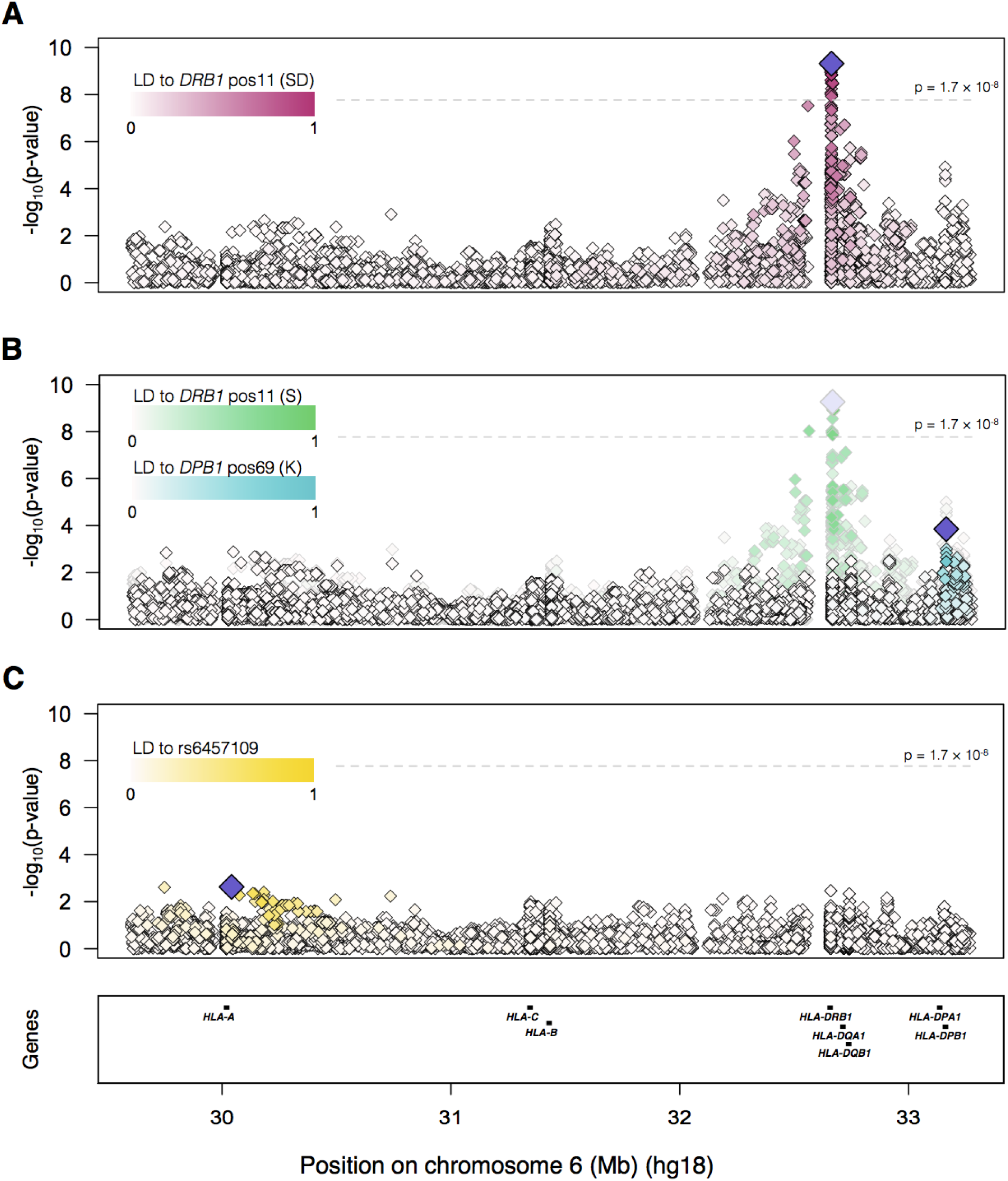
Association and conditional testing in *HLA-DRB1*. **A.** Initial association testing in the MHC revealed a genome-wide significant signal at *HLA-DRB1* position 11 (presence of serine (S) or aspartic acid (D), purple diamond). **B.** Conditioning on the presence of aspartic acid at position 11 left the association signal essentially unchanged (green and white diamonds, grey outline); presence of serine remained the strongest association (pale purple diamond). Conditioning on the presence of serine at position 11 (aquamarine and white diamonds, black outline), dramatically mitigated the association signal, indicating that presence of serine explains the bulk of the association at *DRB1* position 11. Presence of lysine (K) at position 69 in *HLA-DPB1* remained modestly associated (dark purple diamond). **C.** Conditioning on lysine at position 69 in *HLA-DPB1* removes the remainder of the signal (rs6457109, dark purple diamond, p = 0.0024).

Serine at position 11 in *HLA-DRB1* is positioned in the middle of what is known as the YST- motif^35,36^ in *HLA-DRB1*; the motif is comprised of tyrosine (Y) at position 10, serine (S) at position 11, and threonine (T) at position 12. All three amino acids are in perfect linkage disequilibrium (LD) with one another (i.e., LD = 1 between all residue pairs). Additionally, the residues in the YST-motif are also in perfect LD with a fourth residue, presence of serine (S) or glycine (G) at position 13 in *HLA-DRB1* (OR = 2.47 [95% CI: 1.84 - 3.32, p = 1.44×10^−9^ in the MHC analysis). Thus, all four amino acid configurations show statistically identical association to JIA-associated uveitis (Table 2). To further decipher the specific residue(s) driving the association in uveitis, we performed a series of likelihood ratio tests. In brief, a likelihood ratio test compares the fit of two models to the observed data and indicates which of the models, if either, best fits the data. We found that the S (or Y or T) amino acids best fit the data (p = 1.48 × 10^−10^, compared to a model containing only PCs and sex as independent variables). Including the S/G residue at position 13 as well as the YST motif only modestly improved the fit of the model (likelihood ratio test p = 0.043, compared to a model containing serine at position 11 only); neither the S nor G residues alone at position 13 improved the model (p = 0.70 for both tests).

A recent MHC fine-mapping study of the JIA categories tested here mapped the primary disease association signal to the G residue at position 13 in *HLA-DRB1*, and additionally identified an independent risk for serine at the same position. Analyzing all JIA patients (with and without uveitis), we also observed an association at glycine-13 (**Supplementary Table 3**). We sought to disentangle the previously-described association signal in all JIA (residues S or G at position 13) from the JIA-associated uveitis signal (residue S at position 11) observed here. We adjusted for either G or S at position 13 in the JIA-associated uveitis vs. non-uveitis JIA analysis and found the signal at serine-11 to be only moderately affected (serine-11 adjusted for glycine-13: OR = 2.41 [95% CI: 1.75 - 3.32], p = 6.49×10^−8^; adjusted for serine-13; OR = 2.61 [95% CI: 1.74 - 3.93], p = 3.83×10^−6^). These results indicate that the association signal in uveitis at serine-11 (or tyrosine-10 or threonine-12) cannot be fully explained by the JIA-associated amino acids in position 13 in *HLA-DRB1*.

### Sexual dimorphism in JIA-associated uveitis

Since uveitis is epidemiologically known to be more prevalent in females with oligoarticular JIA,^37^ we wanted to formally test if the primary signal in uveitis showed evidence of sexual dimorphism in our sample. In testing female samples only, we found a genome-wide significant signal at serine-11 in *HLA-DRB1* (OR = 3.30 [95% CI: 2.26 - 5.83], p = 7.61×10^−10^); we found no evidence for such an association in the male samples (OR = 1.41 [95% CI: 0.86 - 2.31], p = 0.177; Table 2 **and Supplementary Figure 5**). Although the most significant association with uveitis in females was mapped to amino acid position 233 in the cytoplasmic domain of *HLA-DRB1* (presence of threonine (T), OR = 0.30 [95% CI: 0.20 - 0.41], p = 3.50×10^−10^; Table 2), this position is in nearly perfect LD (r^2^=0.98) with tyrosine (Y) at position 10, serine (S) at position 11, threonine (T) at position 12 (the YST-motif), and position 13 in *HLA-DRB1* (**Supplementary Figure 6**), and consequently yield a nearly identical association (Table 2).

To further explore the potential for a sex-specific effect at *HLA-DRB1* position 11, we ran the association test at serine-11 across all samples, including an interaction term between sex and the imputed dosage at serine-11. We found a significant effect of the interaction term (OR = 2.25 [95% CI: 1.22-4.15] p = 0.0096), indicating that the serine-11 signal in *HLA-DRB1* was indeed a sex-specific effect.

### In Silico Peptide Binding to HLA-DRβ 1

Polymorphisms in the beta chain specify the peptide binding preference of the HLA-DR molecule. Serine at position 11, as well as the adjacent associated amino acids at position 10 - 13, are located in the bottom of the antigen-binding groove of the HLA-DRB1 protein (Figure 2), suggesting that different peptide-binding preferences of HLA-DRB1 may confer risk for developing uveitis. To explore if the presence of serine at position 11 affects peptide-MHCII interactions, we compared the predicted binding affinity for 13 common HLA- DRB1 allotypes (representing 79% of *DRB1* alleles in cases) using a large panel of >80,000 peptides based on human iris proteome data (**Materials and Methods** and **Supplementary Table 4**) using the *NetMHCIIpan* server.^38^ Briefly, the neural network-based *NetMHCIIpan* algorithm is capable of reliably detecting differences between peptide-binding repertoires of highly similar MHC class II molecules.

**Figure 2.**
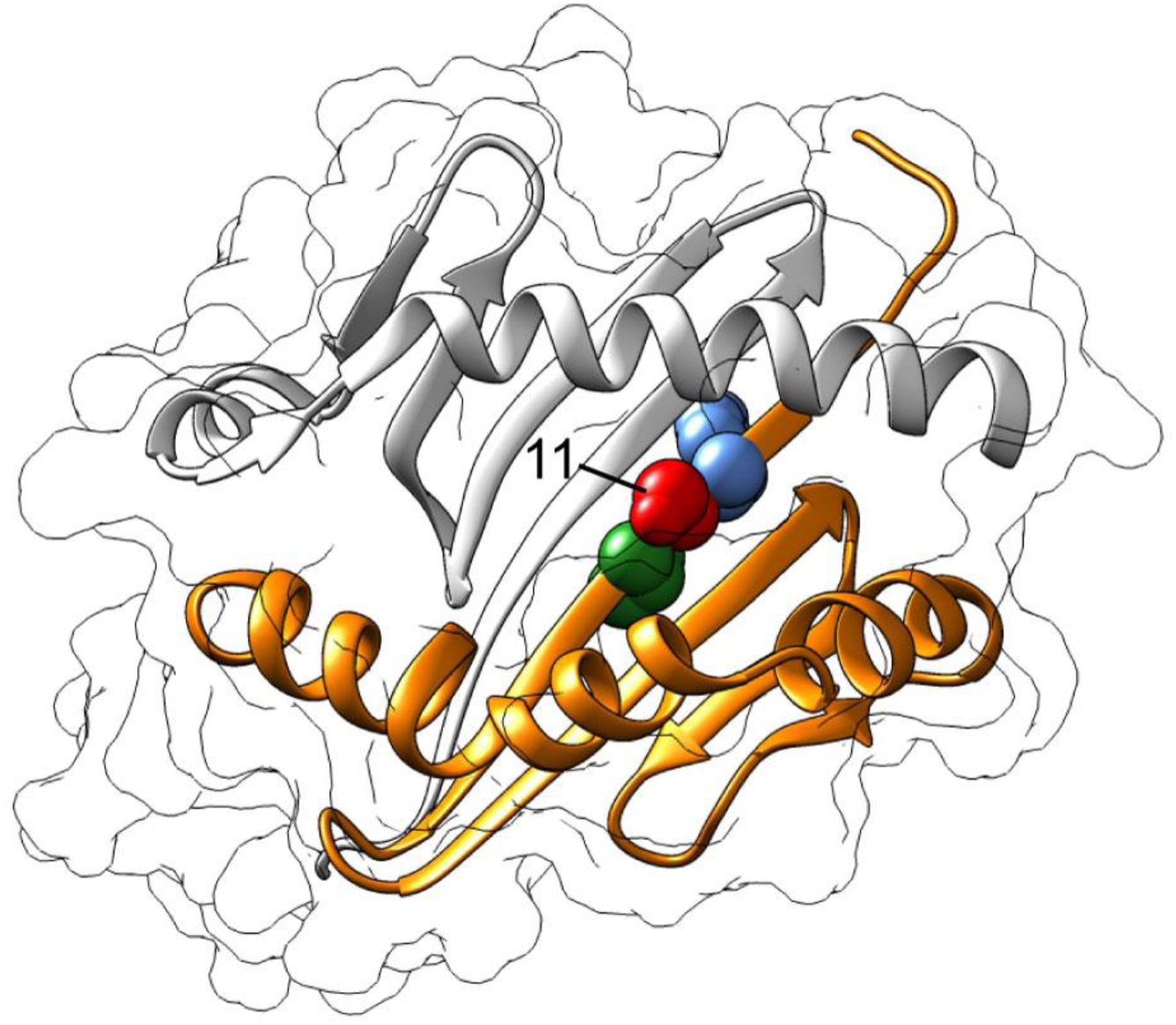
Three-dimensional ribbon model for HLA-DR (Protein Data Bank entry: 3pdo). The molecule is positioned to provide a view from the top of the peptide-binding groove. The beta-chain (DRß) is highlighted in orange. Amino acid serine at position 11 (red) is located in the bottom center of the peptide-binding groove of HLA-DRß1. Adjacent amino acid tyrosine at position 10 (green) and threonine at position 12 (blue) and serine at position 11 are displayed as spheres. 3D structure was produced using UCSF Chimera.^83^

To identify *HLA-DRB1* alleles that encode proteins with similar predicted binding preferences, we performed unsupervised hierarchical clustering. Clustering of the peptide affinity data discerned two major clusters of classical alleles strikingly similar to the distribution of serine at position 11 (**Supplementary Figure 7**). We observed that the average binding affinity of the peptide panel was higher for the *HLA-DRB1* alleles that encode serine at position 11 versus alleles that have other amino acids at this position (Wilcoxon signed-rank test, p = 3.44×10^−136^, **Supplementary Table 5**). To compare these two clusters, we then computed the ratio of the average binding affinity of the 6 HLA-DRB1 allotypes that contain serine-11 and the 7 that have other amino acids at this position (**Supplementary Table 5**). Since MHC class II molecules at the cell surface present repertoires that are skewed in favor of high affinity binders,^39^ we selected for peptides with an (IC_50_) affinities < 500 nM (an affinity of < 500 nM is routinely used as a threshold for potential immunogenicity) or < 50nM (strong binding peptides) for HLA-DRB1 allotypes with serine-11 (**Materials and Methods**). The data indicated that peptides with an intermediate or high affinity for allotypes containing serine-11 have less affinity to allotypes that contain other amino acids at position 11 (**Supplementary Table 5**).

### Previously-described associations to JIA and uveitis

A number of HLA types and amino acid positions have been demonstrated to be associated to JIA, uveitis and related autoimmune diseases through genome-wide association and candidate gene studies.^20,40,41^ We looked up 10 previously-associated HLA types and amino acids in our GWAS of (1) JIA and unaffected controls and (2) uveitis and unaffected controls. Seven of the 10 associations appeared in our data, and all associations were nominally significant (p < 0.05; **Supplementary Table 6** and **Supplementary Figure 8**).

## Discussion

Through interrogation of imputed HLA types and amino acids, we have identified the amino acid serine at position 11 in the *HLA-DRB1* gene as strongly associated to increased risk of uveitis in female patients with JIA. This association indicates that, though JIA with uveitis and JIA without uveitis share clinical features and genetic risk factors (and in particular, HLA-specific genetic risk factors^20^), there is genetic architecture specific to uveitis.^4,42,43^ Specifically, we use conditional testing and likelihood ratio tests to show that the previously reported signals in JIA at position 13 is a distinct signal from the uveitis signal at position 11 in *HLA-DRB1*. These findings suggest that the two phenotypes are likely to have biological features and progressions that are partially distinct.^44^

Association testing initially revealed either serine or aspartic acid at position 11 in *HLA- DRB1* as the most-associated variant, a signal we determined was primarily driven by serine. However, the perfect linkage disequilibrium between serine, tyrosine (position 10), and threonine (position 12) makes it impossible to disentangle the three amino acids in a statistical framework; a subset of the amino acids, or all three (that is, the YST-motif) may be key to disease onset. Importantly, though all three residues are located at the bottom of the HLA-DRB1 peptide-binding groove, only serine-11 is positioned towards, and, thus most likely interacting with, binding peptide epitopes (Figure 2). Previous work has identified serine-11 as the strongest risk factor in seronegative rheumatoid arthritis (RA). Some JIA patients, including those with uveitis, might be categorized as seronegative RA by the time they reach adulthood, and seronegative RA is considered to genetically mirror the uveitis-prone JIA categories.^20^ In contrast, serine-11 is highly protective against seropositive RA, a biologically distinct form of RA in which uveitis is not common (<1% of cases).^45,46^

Sex-specific analyses in the MHC further revealed the *HLA-DRB1* signal was unique to female samples: the interaction of sex and the *HLA-DRB1* serine-11 genotype was significant, and the association signal at serine-11 was entirely absent when the analysis was restricted to male samples only. The reduced number of cases (N = 56 male JIA-uveitis cases and N = 102 male non-uveitis JIA controls) and therefore reduced power in the male-only analysis may explain the absence of a signal at serine-11. However, we had 98.5% power to detect an association of p < 0.05 at serine-11 in the male-only analysis, assuming the odds ratio in males was the same as in females (OR = 3.30). Assuming a more modest effect in males (OR = 2.00), we still had 75.2% power to detect a signal at p < 0.05. To detect potentially more subtle effects in males (OR < 2), larger case collections will be necessary to improve power. Nonetheless, the interaction test of sex and serine-11 provides compelling evidence for a sexually-dimorphic signal in *HLA-DRB1*.

In-depth data on sex-specific epidemiology of uveitis associated with JIA are sparse, however, the typical bilateral chronic anterior uveitis has been reported to occur more frequently in females.^37,47^ This genotype-by-sex interaction informs an exciting field of future research that aims to elucidate how the uveitis risk mechanisms underlying the *HLA- DRB1* signal may be mediated by sex-specific factors, such as hormone regulation. Further, the sex-specific pattern of genetic association may help to resolve previously reported evidence of sexual dimorphism with respect to severity or disease course in children with JIA-associated uveitis.^48^ Lastly, the finding indicates that sex stratification may be beneficial for future clinical trials, as some therapeutic agents may be more efficacious in females or males only.^49,50^

HLA-DRβ 1, like other MHC class II molecules, is essential to immunity and orchestrates downstream immune responses by presenting a dynamic cargo of thousands of different peptides for scrutiny by T helper cells.^51^ Both JIA and uveitis are considered diseases mediated by T helper cell subsets.^4^ The strong association of an amino acid motif (tagged by serine-11) in *HLA-DRB1* suggests that predetermined peptide preferences may affect the regulation of T helper cells. To investigate the immunological impact of the motif in *HLA-DRB1*, we used a “reverse immunology” approach^38^ (**Materials and Methods**), performed using data from proteins experimentally observed in human iris.^52^ Since JIA-uveitis patients frequently have circulating ANAs directed to iris tissues, we specifically focused on nuclear proteins to provide a disease-relevant dataset for modelling. Although the large panel of putative antigens that we tested was by no means exhaustive (**Supplementary Table 4**), the experiment demonstrated that the presence of a serine-11 motif in *HLA-DRB1* is accompanied by distinguished changes in binding affinity in the resulting protein (**Supplementary Table 5, Supplementary Figures 7**), suggesting that antigen presentation by the HLA-DRβ1 protein may underpin the uveitis susceptibility. Risk alleles of *HLA-DRB1* that encode serine-11 (e.g. *HLA-DRB1*11*, **Supplementary Table 2**) may communicate distinct ‘peptidomes’ that influence T helper cells and significantly increase the likelihood of downstream immune responses (e.g., ANA) to the eye. Further, ANA-positivity is modestly correlated with presence of serine-11 in the cases studied here (Pearson’s r= 0.26, *P =* 4.9×10^−4^). Functional studies using HLA-proteomic approaches will be necessary to experimentally validate and systematically dissect the complex downstream effects of serine-11 on tissue-specific antigen presentation by HLA-DRβ1 in uveitis.

The relatively high frequency of serine-11 in the JIA cases that did not develop uveitis (Table 2) indicates the likely involvement of additional (epi)genetic and environmental factors. Indeed, genes outside the MHC have been implicated in JIA-uveitis, including a polymorphism in *VTCN1*^53^ and variants near the immune genes *TRAF1* and *C5*.^54^ More than half of patients with JIA-uveitis display elevated titers of anti–parvovirus B19 antibody in ocular fluids,^43,55^ and human parvovirus B19 infection can cause various clinical features, including arthritis, uveitis and ANA production.^56,57^ In addition, immunity to Parvo-B19 infection is strongly associated with *HLA-DRB1* and involves T helper cells. These observations collectively outline a complex interplay of both biologic and environmental factors in JIA-uveitis.^58–61^

Though we were able to identify the serine-11 association in this collection of JIA-uveitis and JIA without uveitis samples, our analyses leave unanswered questions as to the genetic underpinnings of uveitis, and potential sex-based differences in their pathogenesis. Given our sample size, we were well powered to detect common (frequency >10%) and highly-penetrant (odds ratio > 3, **Supplementary Figure 9**) variation that associates with increased uveitis risk compared to JIA individuals without uveitis. However, we are underpowered to identify common, modestly-penetrant variation - the hallmark genetic feature of complex phenotypes^62,63^ - that may further distinguish the two phenotypes. In the current study, we included only patients for whom we were able to obtain detailed ophthalmological follow-up for ≥ 4 year (±90% of uveitis develops within 4 years after onset).^64^ While imperative to revealing the serine-11 association, this strategy limited case ascertainment. Future genetic studies will likely benefit from broader inclusion of cases, including essential, detailed phenotype information,^7,65^ in order to identify additional genetic associations that are shared across or distinct to particular uveitis subtypes of gender.^66^ In addition, clinical studies - including clinical trials - may be improved by being well-powered in both sexes to enable sex-specific investigations.

While much progress has been made in understanding the biology of JIA,^16,20^ our understanding of the genetic landscape of uveitis is more limited. Only a handful of candidate gene studies^41^ and HLA-specific analyses^4,17–19^ have investigated uveitis as a phenotype separate from all JIA. A recent GWAS identified *HLA-DRB1*1501* as a risk factor for uveitis,^67^ and recent studies in JIA-associated uveitis point toward a role for infiltrated plasma cells^68^, T helper cells^4^, and changes in the ocular fluid microenvironment.^43,69,70^ Future, larger genetic studies comparing JIA-uveitis and JIA without uveitis, with and without sex-specific analysis, will likely expand our understanding of uveitis pathogenesis in a number of ways. Broad case inclusion will elucidate whether subtype-specific architectures exist and larger samples will improve statistical power to identify more modest-effect disease-modifying variants both within and outside of the MHC. For example, this may substantiate the association of additional HLA alleles (such as *HLA-DPB1*) with uveitis in JIA (Figure 1).

Larger studies may also help in revealing clinically-useful biomarkers. Risk factors such as young age and presence of ANA, assumed to reflect aberrant immune activation,^37,40,64,71^ indicate patients potentially at higher risk of uveitis but are in no way predictive of disease outcome. An easily-testable biomarker such as a genotype would be highly impactful in the management of uveitis, as it would help in overcoming the challenge of detecting the insidious onset of disease^7,48,65^. Intriguingly, after deeper examination of the phenotypic data, we found that 99% of female uveitis cases in our study carry at least one copy of the variant coding for serine-11. The only female uveitis case who did not carry serine-11 in *HLA-DRB1* appeared to suffer from ANA-negative oligoarthritis with mild vitritis and peripheral multifocal choroiditis in the absence of anterior segment inflammation; this ocular finding is atypical for JIA, and thus, according to our inclusion criteria, this sample was likely improperly included at cohort collection. Prospective studies in larger populations, including detailed clinical evaluation of development of uveitis and secondary uveitis phenotypes, will be necessary to dissect the potential of serine-11 or other genotypes as biomarkers for disease risk. Regardless, the current study justifies further genetic analysis as a potentially powerful instrument in moving analysis of the genetic architecture of uveitis towards accurate and efficient clinical decision tools.

The results of our study represent a key step in understanding the pathogenesis of uveitis in JIA, helping to discern biologically shared and distinct features of JIA-uveitis and JIA without uveitis. Future work will allow us to further disentangle the two phenotypes and evaluate shared and distinct disease etiology. Unraveling the pathogenesis of the two diseases will improve understanding of the key pathways that trigger disease. By pinpointing and understanding these mechanisms, we can potentially identify clinically-useful biomarkers that stratify patients for disease risk in order to improve personalized care, and catalyze future lines of research in precision medicine. Collectively, this work will advance our progress towards treating and preventing sight-threatening complications of uveitis in children with JIA.

## Material and methods

### Patient collection

#### JIA and uveitis samples

A total of 576 patients with oligoarticular extended, oligoarticular persistent or rheumatoid factor negative polyarticular JIA, of whom 214 (37%) had developed uveitis, were enrolled in this study. These patients were collected from two independent cohorts: (1) a Dutch and cohort, and (2) a cohort comprised of samples collected in Germany, Belgium and Switzerland. For simplicity, we will refer to these as the Phase 1 and Phase 2 sample sets, respectively. Three-hundred eighty four patients were included in the Phase 1 samples (137 uveitis samples) and 192 patients were included in the Phase 2 samples (77 uveitis samples).

JIA was diagnosed according to the criteria of the International League of Associations for Rheumatology (ILAR), or by former criteria (e.g. European League Against Rheumatism (EULAR)).^2,72^ All patients were screened by an ophthalmologist according to the guidelines of the Academy of Pediatrics and patients with no clinical signs of uveitis had an ophthalmologic follow-up of at least 4 years after onset of JIA.^73^ Patients with at least trace cells or more in the anterior chamber and treated with at least topical steroids during ophthalmologic examinations, were diagnosed with JIA-associated (anterior) uveitis.

DNA material of JIA patients with and without uveitis from Phase 1 were collected at the University Medical Center Utrecht, University Medical Center Leiden, Erasmus Medical Center Rotterdam, Academic Medical Center Amsterdam and Radboud University Medical Center Nijmegen (all based in the Netherlands). Samples from Phase 2 were collected within the ICON study and provided by the ICON biobank at the Westfälische Wilhelms-Universität Münster (ICON-JIA Study, Germany), the University Hospitals Leuven (Belgium), the University Children's Hospital at Zurich (Switzerland).

#### Population-level control samples

Genotype data from 394 unrelated and unaffected Dutch samples were used as population controls and had been previously genotyped using the same platform as the JIA and uveitis samples contained in this study.^74^

### Data collection

Information such as JIA subtype, sex and uveitis status were collected from all patients; presence of antinuclear antibodies (ANA) could be collected from the majority of patients (68%) from samples included in Phase 1 and 2.

This study was approved by the local Institutional Review Boards and is in compliance with Helsinki principles. Informed consent was obtained from all participating patients if they were 18 years or older, from both parents and patients if they were 12-18 years of age, and from parents only if they were younger than 12 years old.

### Genotyping and quality control

All samples were genotyped using the Infinium HumanOmniExpress-24v1.1 and v1.2 arrays, which contain ∼700,000 SNPs. For Phase 1, all JIA cases (with and without uveitis) were genotyped together; unaffected controls were genotyped on the same array, but as a separate batch. All Phase 2 samples were genotyped together.

#### Sample-level quality control

Before performing imputation, we ran sample-level and SNP-level data quality control, following standard genome-wide association quality control steps. All steps, thresholds, and number of removed samples and SNPs are summarized in **Supplementary Table 7**. Briefly, we used PLINK 1.9^31^ to first check sample-level missingness across all samples and removed all samples with missingness >5%. Next, for the remainder of sample-level quality control, we reduced the dataset to only high-quality SNPs: SNPs with minor allele frequency (MAF) > 10%; SNPs outside the MHC (chromosome 6), lactase locus (chromosome 2), and the inversions on chromosomes 8 and 17; SNPs with missingness < 0.1%; and SNPs linkage disequilibrium (LD) pruned at an LD (r^2^) threshold of 0.2.

With this high-quality set of SNPs, we performed principal component analysis (PCA) using EIGENSTRAT.^75^ We defined samples to be of European ancestry if their principal component (PC) 1 and PC 2 values were within 6 standard deviations of the European-ancestry populations included in the HapMap 3 dataset (the CEU population, samples of Northern and Western European ancestry living in Utah; and the TSI population, Toscans living in Italy). In addition, we calculated inbreeding coefficients for all samples and removed samples further than 3 standard deviations from the distribution. Related samples (pi-hat > 0.125) were also removed. Finally, using all SNPs available on chromosome X, we performed a sex check to identify samples with mismatching genotypic and phenotypic sex; samples with mismatching genotype and phenotype information were removed from the analysis.

#### SNP-level quality control

We dropped failing samples from the data and then proceeded to perform SNP QC. All SNPs with missingness >5% were removed, as were SNPs with frequency <1% (too rare to be analyzed in the logistic regression framework, given the sample size). SNPs out of Hardy-Weinberg equilibrium in controls or across the full dataset were removed (p < 1×10^−6^ in Phase 1; p < 1×10^−3^ in Phase 2), as were SNPs with significant differential missingness between cases and controls (p < 5×10^−2^ in Phase 1; p < 1×10^−3^ in Phase 2).

Once sample- and SNP-level QC were complete and the failing samples and SNPs had been removed from the data, we performed a second iteration of PCA to ensure that cases and controls were overlapping in PCA space (i.e., there was no obvious population stratification in the data) (**Supplementary Figure 10**).

### Prephasing and imputation

#### Prephasing

Once sample- and SNP-level were complete, the Phase 1 and Phase 2 data were phased separately, as they were genotyped and QC’d separately, using SHAPEIT2.^23^ As the sample size was >100 samples, prephasing was run without a reference panel, per the SHAPEIT2 recommendations.^23^

##### Genome-wide imputation

Following the prephasing, we imputed all samples using the imputation reference panel constructed through whole-genome sequencing of 2,504 samples in 1000 Genomes Project (Phase 3).^28^ Briefly, the 1000 Genomes Project Phase 3 data is comprised of samples collected from the Americas, Africa, East Asia, Europe and South Asia. Samples were whole-genome sequenced at ∼80x coverage across the exome and at ∼4x coverage in non-exome regions. We imputed the prephased haplotypes using IMPUTE2^24,76^; the data were phased in windows 5 megabases long, with a buffer region of 250kb. We set the effective sample size (Ne) to 20,000, per IMPUTE2 recommendations, and set k_hap to 1,000, as our sample was of European ancestry and the 1000 Genomes Phase 3 imputation reference panel contains ∼1,000 haplotypes from European-ancestry individuals.

##### Imputation of HLA classical alleles and amino acids

To impute amino acids and HLA alleles in the MHC, we used the SNP2HLA pipeline.^29^ In short, the SNP2HLA pipeline uses a reference panel assembled through HLA typing of 5,225 European-ancestry individuals collected by the Type 1 Diabetes Genetics Consortium.^77^ The panel includes SNPs and amino acids in the MHC, as well as HLA types from the Class I (*HLA-A*, *HLA-B*, and *HLA-C*) and Class II (*HLA-DRB1*, *HLA-DQA1*, *HLA-DQB1*, *HLA-DPA1*, and *HLA-DPB1*) HLA genes. HLA types are imputed to 2- and 4-digit resolution. The SNP2HLA pipeline uses BEAGLE^78^ to phase the data and then impute.

Post-imputation, we checked the sum of the dosages across each HLA gene for each individual. Samples with dosages >2.5 at any one of the HLA genes (induced through imprecision in the imputation) were dropped from further analysis.

#### Association testing

##### Genome-wide association testing

To perform genome-wide association testing, we coded JIA samples with uveitis as cases and JIA samples without uveitis as controls. First, we performed a genome-wide association study (GWAS) in Phase 1 and Phase 2 individually, to check the overall behavior of the genome-wide test statistics (lambda_Phase1_ = 1.008, lambda_Phase2_ = 1.017). GWAS were performed using PLINK 1.9^31^ using an additive logistic regression model, correcting for the top two principal components and sex. Data were then meta-analyzed using METAL.^32^ To ensure we were analyzing SNPs with high-quality imputation, we only analyzed common SNPs (MAF > 1%) with imputation quality (info) score > 0.7. In the Phase 1 data, additional GWAS were performed comparing samples with JIA and uveitis to unaffected controls, as well as JIA samples without uveitis to unaffected controls; results were concordant with studies previously performed in these phenotypes (**Supplementary Table 6**).

##### Association testing in the MHC

Similar to the genome-wide association testing, additive logistic regression in the Phase 1 and Phase 2 data was first performed separately to check the overall behavior of the data. Then, the dosages were merged together, and logistic regression was performed on the dosage data using PLINK1.9,^30,31^ correcting for the top 5 principal components, sex, and phase. To ensure that this mega-analysis approach was appropriate, we additionally performed an inverse variance-weighted meta-analysis of the two phases, and found that the results were highly concordant (Pearson’s r of genome-wide betas = 0.95). To identify potential independent signals within the MHC, we performed a follow-up regression by conditioning on the top-most variant (Figure 1).

#### In silico peptide binding prediction to *HLA-DRβ1*

Proteome data from human iris tissues (2,959 nonredundant proteins) was used as a representative source of proteins present in iris tissue.^52^ JIA-uveitis patients commonly have antinuclear antibodies (ANA) directed to iris tissues, thus, we focused on nuclear iris proteins to generate a potentially disease-relevant dataset. Protein accession numbers were extracted to filter in UniProt (Universal Protein Resource) for proteins with a subcellular location annotation limited to the nucleus.^79^ We selected 147 proteins (**Supplementary Table 4**) that fulfilled these criteria and their full length amino acid sequences were fed into the neural network of the *netMHCIIpan3.1* server. The netMHCIIpan algorithm^38^ was selected for its high quality performance in peptide binding prediction for any MHC class II allele for which the sequence is identified, including those for whom only few experimental peptide binding affinities (limited allele-specific training data) are available. We ascertained this by testing the predicted binding affinity of several hundreds of experimentally identified 15-mer peptide ligands eluted from HLA-DRβ1:01,^80,81^ which revealed that >95% were correctly picked up by netMHCIIpan as ligands for DRβ1:01 (IC_50_ < 500nM, data not shown).

Next, we tested the predicted affinities of all 83,686 overlapping 15-mer peptides from the selected 147 proteins in *netMHCIIpan3.1*. that tentatively could be presented by HLA-DR for binding to representative four-digit alleles of *HLA-DRB1* (\**01:01, *03:01, *04:01, *07:01, *08:01, *09:01, *10:01, *11:01, *12:01, *13:01, *14:01, *15:01,* and \**16:01*). These alleles account for 79% of *HLA-DRβ1* alleles in the JIA cases. The affinity data was log-transformed to a value between 0 and 1 using: 1-log(IC_50_nM)/log(50,000).^38^ To categorize *HLA-DRB1* allotypes (alleles) with similar predicted binding preferences, we performed unsupervised hierarchical clustering. Heatmaps were created based on the Euclidean distance measure and the Ward’s linkage method using the MetaboAnalyst server.^82^ We computed the ratio of the average binding affinity of HLA-DRβ 1 molecules that contain Serine-11 in the peptide-binding groove (*\*03:01, *08:01, *11:01, *12:01, *13:01,* and *\*14:01*) over the average binding affinity of HLA-DRβ1 proteins that have other amino acids at this position (\**01:01, *04:01, *07:01, *09:01, *10:01, *15:01,* and \**16:01*) as a measure for the overall difference in predicted binding affinity for each peptide. A frequency distribution of the ratio of the binding affinities was plotted for the entire set of peptides (including low and nonbinding peptides), potential ligands for DRβ1 with a binding affinity stronger than half maximal inhibitory concentration (IC_50_) of 500 nM, or peptides with high binding affinity for Serine-11 encoding alleles(IC_50_< 50nM).

## Acknowledgements

The following ICON-JIA study group collaborators have contributed to the DNA samples: Universitätsmedizin Charité Berlin, Tilmann Kallinich; Medizinische Hochschule Hannover, Angelika Thon; Universität Tübingen, Jasmin Kümmerle-Deschner; Prof.-Hess-Kinderklinik Bremen, Hans-Iko Huppertz; Asklepios Kinderklinik Sankt Augustin, Gerd Horneff; Olgahospital Stuttgart, Anton Hospach; Kinderkrankenhaus der Stadt Köln, Kirsten Mönkemöller; Deutsches Zentrum für Kinder- und Jugendrheumatologie Garmisch-Partenkirchen, Johannes-Peter Haas; St. Joseph-Stift Sendenhorst, Gerd Ganser; Kinderrheumatologische Praxis am AK Eilbek Hamburg, Ivan Foeldvari; Deutsches Rheuma-Forschungszentrum Berlin, Jens Klotsche.

## Contributors

JJWK, JHB, and SLP led the study. AJWH, SLP and JJWK wrote the paper. SLP, AJWH, and JJWK performed the data and statistical analysis. JHB, RC, MWS, TRDJR, SSMK, PCEHM, AJWH, CW, TS, DF, RAO, TRDJR, AIDH, SH, NSD, EPAHH, MAJR, and NW contributed to the patient ascertainment, sample collection and/or genotyping. All authors reviewed the final manuscript.

